# Enhanced inference of ecological networks by parameterizing ensembles of population dynamics models constrained with prior knowledge

**DOI:** 10.1101/686402

**Authors:** Chen Liao, Joao B. Xavier, Zhenduo Zhu

**Affiliations:** Program for Computational and Systems Biology, Memorial Sloan-Kettering Cancer Center, New York, NY, United States of America; Department of Civil, Structural and Environmental Engineering, University at Buffalo, Buffalo, NY 14260, United States of America

**Keywords:** Lotka-Volterra model, time-series data, summary food web, ecological network inference, ensemble method, invasive species

## Abstract

**Background:** Accurate network models of species interaction could be used to predict population dynamics and be applied to manage real world ecosystems. Most relevant models are nonlinear, however, and data available from real world ecosystems are too noisy and sparsely sampled for common inference approaches. Here we improved the inference of generalized Lotka-Volterra (gLV) ecological networks by using a new optimization algorithm to constrain parameter signs with prior knowledge and a perturbation-based ensemble method.

**Results:** We applied the new inference to long-term species abundance data from the freshwater fish community in the Illinois River, United States. We constructed an ensemble of 668 gLV models that explained 79% of the data on average. The models indicated (at a 70% level of confidence) a strong positive interaction from emerald shiner (*Notropis atherinoides*) to channel catfish (*Ictalurus punctatus*), which we could validate using data from a nearby observation site, and predicted that the relative abundances of most fish species will continue to fluctuate temporally and concordantly in the near future. The network shows that the invasive silver carp (*Hypophthalmichthys molitrix*) has much stronger impacts on native predators than on prey, supporting the notion that the invader perturbs the native food chain by replacing the diets of predators.

**Conclusions:** Ensemble approaches constrained by prior knowledge can improve inference and produce networks from noisy and sparsely sampled time series data to fill knowledge gaps on real world ecosystems. Such network models could aid efforts to conserve ecosystems such as the Illinois River, which is threatened by the invasion of the silver carp.

## Background

The study of ecosystems seeks to understand and predict the changes in species composition, dynamics and stability. Pioneered by Robert May [1], ecological network theory proposed that species interactions can be quantified by numerical matrices and be used to study relevant ecosystem properties [2]. Applications to real world ecosystems, however, have remained limited because quantifying species interactions requires laborious field work in well controlled environments [3]. Computational methods that seek to infer ecological networks from laboratory or field data include parameter-free correlation-based algorithms such as Pearson’s correlation coefficients [4], parametric or non-parametric statistical and machine-learning methods such as Bayesian networks [4, 5], non-parametric approaches based on nonlinear state space reconstruction such as the convergent cross mapping [6], and nonlinear parametric models of population dynamics such as Ecopath with Ecosim [7]. Some approaches have been successfully applied to discretized co-occurrence (presence-absence) data [4, 5, 8–10] but inference from continuous time-series data has lagged behind [6].

Multispecies population dynamics models, particularly the generalized Lotka-Volterra (gLV) model (Equation (1)), provide a flexible way to model and link species interactions to their temporal abundance changes. By constructing a gLV model, the underlying ecology is phenomenologically summarized with minimal parameterization: the biological growth is modelled by an exponential growth rate and the fitness effect of each one-way interaction is quantified by a single coefficient with magnitude and sign representing the interaction strength and type respectively. GLV models have been extensively used in theoretical and computational ecology, particularly in studies of microbial communities [11–18], due to their simplicity, tractability, and transparent logic. For example, inferring microbial ecological networks from gut microbiome time series data has revealed a native gut bacterial species that prevents invasion by a pathogenic species [17].

Despite the popularity of gLV to infer ecological networks in microbial ecosystems, its use for macroscopic ecosystems remains limited. The present interest in the human microbiome has produced abundant datasets for microbial ecology. Macroscopic ecological field data, when they are available, tend to be noisy, sparsely sampled and lack replicates [19]. GLV inference (despite many follow-up efforts [12, 20, 21]) is most commonly parameterized by linear regression (LR) [11]: the gLV model is first discretized and transformed into a system of linear equations and then fit by a regularized multilinear regression (see Methods). The numerical discretization of differential equations is significantly error-prone because the calculation of the gradients of noisy data (***g*** in Equation (6)) amplifies and propagates the error forward. Therefore, even the optimal solution to the transformed linear problem can produce a network that recreates the observed dynamics poorly [14]. Moreover, even the signs of inferred interactions may be inconsistent with prior knowledge of food webs whose trophic organization constrains the types of interactions among species in the web. Finally, uncertainty of data can be translated into uncertainty of the single “best” model, making it unreliable to draw scientific conclusions solely based on model without knowing the uncertainty of its associated parameters.

Here we tackled these challenges by developing independent solutions and combining them into one approach to infer the network of species interactions from time-series data of Illinois River fish community. The data has been annually sampled by the Long Term Resource Monitoring Program in the Upper Mississippi River System [22], one of the very few ongoing long-term monitoring programs in large rivers in the United States [23]. Briefly, we introduced a novel optimization algorithm that allows for estimation of the gradients in addition to model parameters. During the optimization, the signs of gLV parameters were constrained based on a summary food web that represents all potential interactions among fish species. By searching the parameter space, we constructed an ensemble of models that harbor distinct sets of parameters but fit the data almost equally well. Using ensemble mean and variance, we were able to make robust inferences/predictions of network structure and dynamics as well as to assess whether or not these network properties are well constrained by the data. Finally, we used the ensemble of models to assess the impact of the silver carp (*Hypophthalmichthys molitrix*), an invasive species in the Mississippi and Illinois Rivers [24, 25] that presents a major problem that may percolate to the Laurentian Great Lakes in the future [26].

## Results

### Fish community varies in space and time

The Illinois River is a major tributary of the Upper Mississippi River, where the long-term monitoring efforts of the fish community spread across six field stations since 1993 (Fig. 1a). To visualize how the fish community structure varied across time and space we first standardized catch-per-unit-effort data to combine fish numbers obtained from the different fishing gears employed (see Methods, Additional file 1: Fig. S1). Then we carried out a principle component analysis (PCA) using data from the normalized abundances of 153 fish species for each year and site (Fig. 1b). The data from each site occupied distinct regions of the PCA plot, indicating distinct fish ecologies in space. The communities, despite regional differences, were most similar between proximal sites. The first component, which explains 12% of the variance in the data, is strongly determined by variations in the common carp and bluegill, two species highly abundant in the Mississippi River upstream from the confluence with the Illinois River (Pool 4, Pool 8, and Pool 13) but less abundant in the Illinois River (LG) and the Mississippi River downstream from the confluence (Pool 26 and OR).

**Fig. 1.**
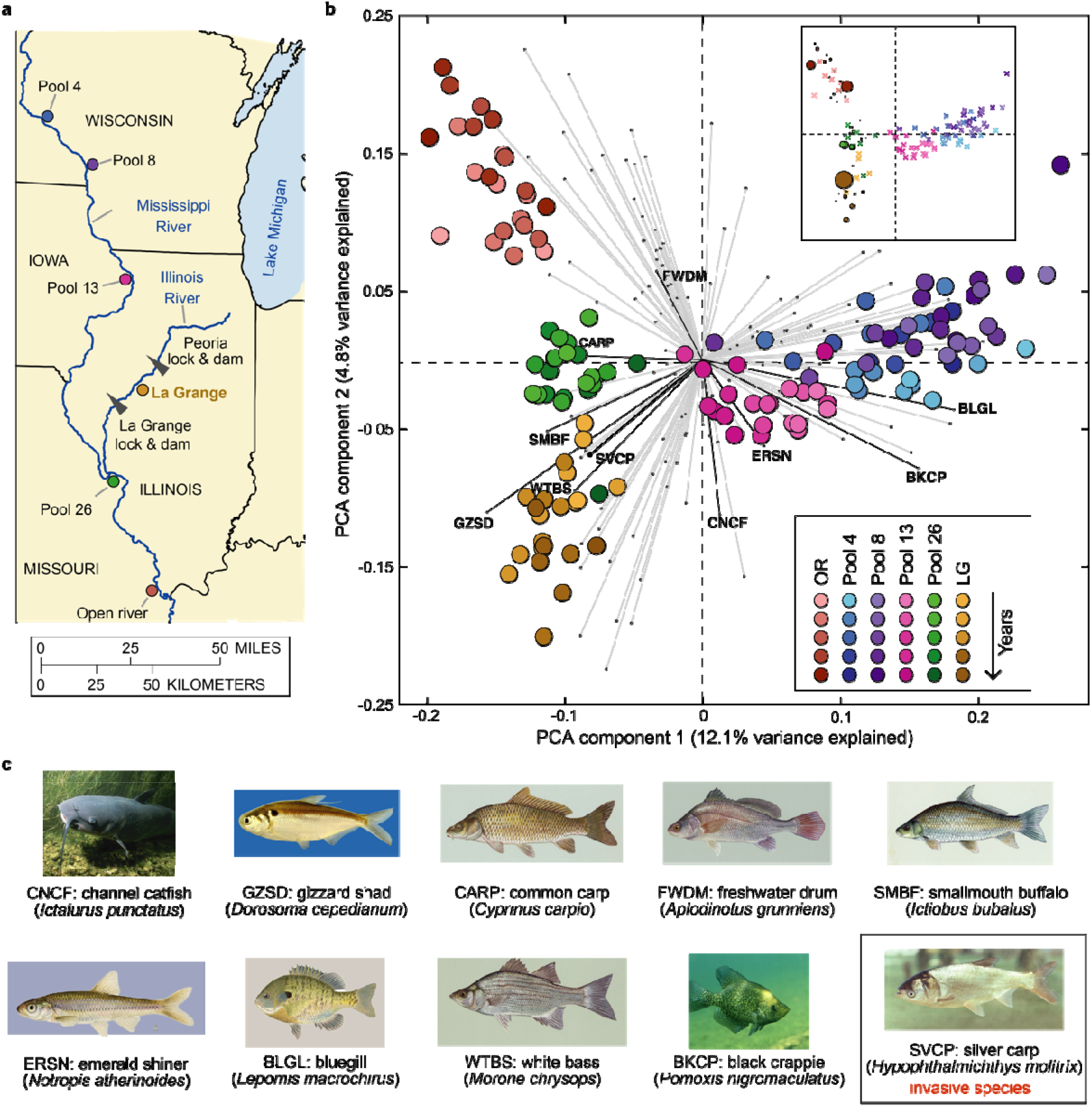
Field measurement provides population dynamics data on the freshwater fish community in the Upper Mississippi and Illinois Rivers. **a** Geographical location of the six stations monitored by the Long Term Resource Monitoring Program. The La Grange (LG) pool, located in the Illinois River, is the focus of the study. **b** Biplot of principle component analysis (PCA). Each circle (“score”) represents the species abundance distribution of fish community associated with a site and year combination. The color brightness of circles indicates the passage of time (from 1993 to 2015): lighter colors represent earlier data. Each line (“loading vector”) represents contribution of an explanatory variable (fish species) to the variance of the first two principle components. For all loading vectors, the top 9 dominant native fish species in the LG pool plus silver carp, an invasive species, are colored in black while all others are colored in light gray. The inset is the same PCA score plot, but the circle size is scaled to be proportional to the abundance of invasive silver carp (samples missing silver carp are represented with crosses). **c** Common names, abbreviations, and species names of the 10 fish species investigated in our study. Fish images were obtained through public domain resources except for silver carp licensed by CC BY 3.0 and gizzard shad provided by Chad Thomas pf Texas State University.

Our PCA illustrates that silver carp (Fig. 1c), one of the four species of invasive Asian carps, has established the lower and middle Mississippi river. The impact of the silver carp was detected in three sites (OR, Pool 26, and LG) over the course of the invasion (Fig. 1b, inset). The Illinois River is known to have one of the highest silver carp densities worldwide [27]. The large silver carp density is obvious in the PCA, which shows that the loading vector for the silver carp aligns well with the La Grange community data (Fig. 1b, in brown). In contrast, the Mississippi sites upstream of the confluence with the Illinois River (Pool 4, Pool 8, and Pool 13) where silver carp are barely found (Fig. 1b, inset) are misaligned with the silver carp vector. Fig. 1b and its inset also reveal the invasion path: silver carp entered the Illinois River at the confluence, rather than continuing to migrate up the Mississippi River. There is grave concern that the invader may enter Lake Michigan through the Illinois River, threatening the Great Lakes’ ecosystems and multi-billion-dollar fishing industry [26].

Among the six observation sites, we focused mainly on the fish community in the LG pool, the only monitoring site along the Illinois River, for two reasons: (1) the pool has both upstream and downstream dams (Fig. 1a) and likely resembles a closed ecosystem that is minimally influenced by immigration and emigration of fish species; (2) the pool has a large population of silver carp (Fig. 1b, inset) and thus can be used to study the impact of this invasive species on the native fish. We chose to model the top 10 most abundant fish species (Fig. 1c, Additional file 2: Table S1)—including 9 native species and 1 invasive species (silver carp)—that together account for 87.1% of the total abundance (Additional file 1: Fig. S2). The ecological effects of the remaining low-abundance species were assumed negligible; we chose not to group these species into one superspecies virtual group to avoid spurious links between that virtual group and the abundant species [14].

### A latent gradient regression algorithm improves gLV parameterization

To reduce the error in numerical approximation of the gradients, we treated the time gradients as latent parameters (their large uncertainty essentially makes them unobserved quantities) and iteratively learned by minimizing error between observed data and model predictions (see Methods, Fig. 2a). We first benchmarked the latent gradient regression (LGR) algorithm by using synthetic data produced by a 3-species gLV model with known parameter values (see Methods, Fig. 2b). In the absence of noise, we show that LGR outperformed LR in data fitting (adjusted R^2^: 99% vs. 36%) and recovered the ground-truth model parameter values (adjusted R^2^: 99% vs. 90%) (Fig. 2b). Using the same benchmark model with noise (see Methods), the LGR’s ability to recover known parameter values was slightly compromised, but still outperformed the LR for curve fitting (Fig. 2c). Finally, nonlinear regression also fitted the data poorly (adjusted R^2^: 53%) and was unable to accurately estimate the ground-truth parameter values (adjusted R^2^: 84%) (Additional file 1: Fig. S3). The convergence rate of nonlinear regression was also much slower than LGR (Additional file 1: Fig. S3).

**Fig. 2.**
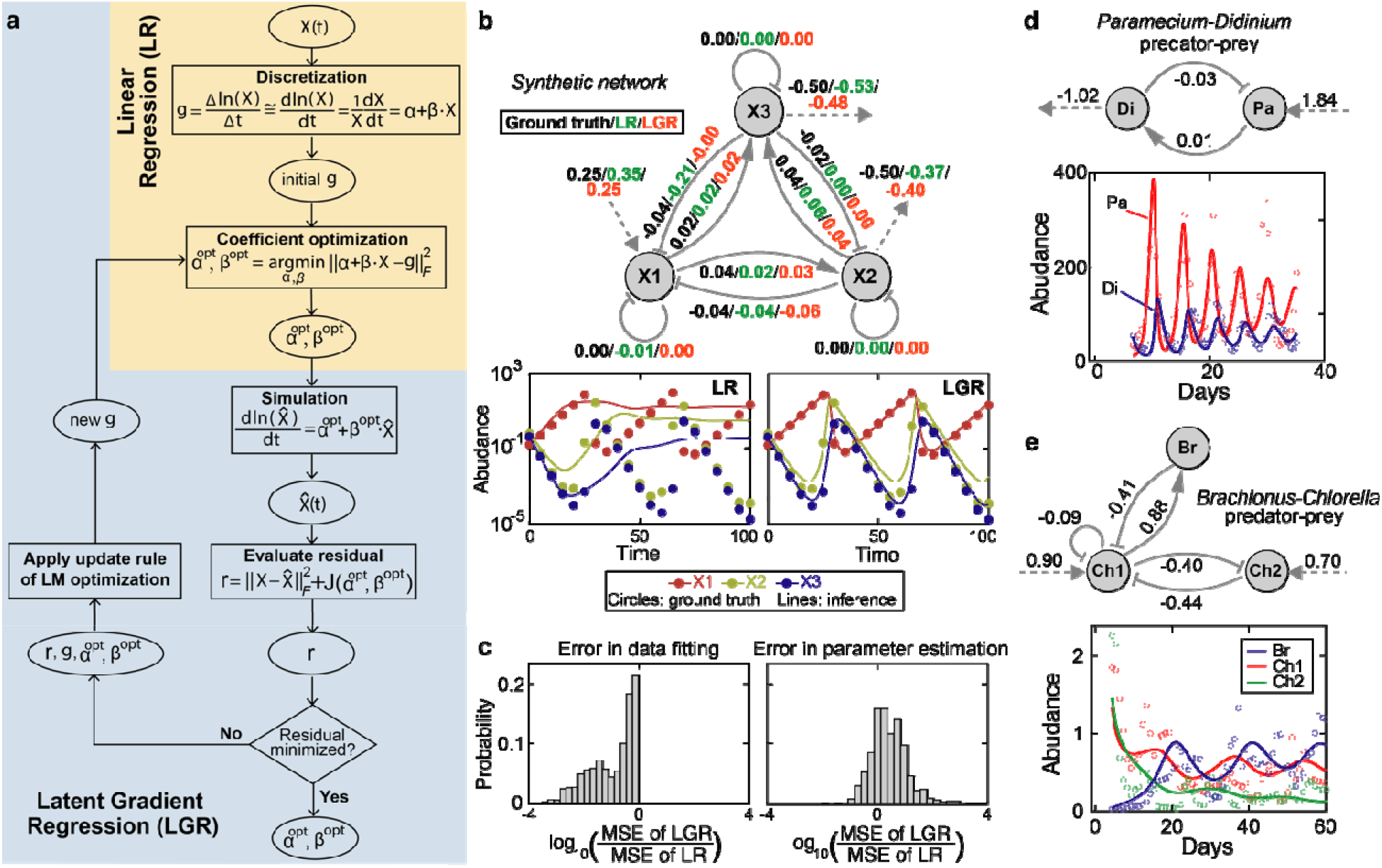
Latent gradient regression algorithm enables parameterization of generalized Lotka-Volterra (gLV) network model. **a** A flowchart showing how linear regression (LR; shaded in light yellow) is expanded to include gradients () as latent parameters in our latent gradient regression (LGR; shaded in light blue) algorithm. : observed time series; : simulated time series; : gLV model coefficients; : gradients (i.e., time-derivatives of ; : penalty function; : Frobenius norm; LM: Levenburg-Marquardt. **b**,**c** Benchmark of the LGR algorithm using synthetic data in the absence (**b**) and presence (**c**) of noise. The synthetic data was generated by a 3-species gLV network model (**b**), where solid arrows represent positive (point end)/negative (blunt end) interactions and dashed arrows represent intrinsic population growth (incoming)/decline (outgoing) in the absence of other species (the same as in **d**,**e**). The best-fit model predictions (lines) are contrasted with the synthetic data (filled circles) in th lower part of **b**. MSE: mean squared error. **d,e** Performance of the LGR algorithm in inferring real ecosystems. **d** The protozoan predator (*Didinium nasutum*)-prey (*Paramecium aurelia*) ecosystem. Unit of abundance in y axis: individuals/mL. **e** The ecosystem of a rotifer predator (*Brachionus calyciflorus*) and two algae prey (*Chlorella vulgaris*). Unit of abundance in y axis: 10 individual females/mL for the rotifer and 10^6^ cells/mL for the algae. In both **d** and **e**, the inferred gLV models are shown in the upper part and their predictions (lines), together with the observed data (empty circles), are shown in the lower part. To eliminate the initial transient period, the first 13 and 4 data points of population dynamics in **d** and **e** were removed respectively.

To test the effectiveness of combining gLV network model and LGR inference algorithm further we analyzed two separate, independently published laboratory predator-prey microbial systems [28, 29], where the interspecific relationships are known and we could use the interaction signs to constrain the inference. GLV inference using LGR successfully identified network structures that reproduced the community dynamics observed experimentally in both datasets (Fig. 2d,e). Quantitatively, the adjusted R^2^ for the two-species *Didinium nasutum*-*Paramecium aurelia* ecosystem and three-species rotifer-algae ecosystems were 74% and 70% respectively. Moreover, the inferred network structure of the rotifer-algae ecosystem agreed with the observed fitness trade-off in survival strategies employed by the two algal clones [29]: the second clone Ch2 grew slower than the first clone Ch1 (the inferred growth rates of Ch1 and Ch2 are 0.9 and 0.7 respectively) but developed resistance to rotifer’s predation (the inferred predation strength of the rotifer on Ch1 and Ch2 are −0.41 and 0 respectively).

### A summary food web of fish community constrains gLV parameters

Food webs that describe trophic positions of prey and predators constrain the signs of interactions between species. We sought to reconstruct a summary food web consisting of all potential interactions among the 10 selected fish species and transform them into parameter sign constraints. Using the summary food web to constrain gLV parameters enables integration of prior knowledge in the network inference process, which not only improves efficiency in searching high-dimensional parameter space but guarantees qualitative agreement between the inferred network and literature data.

As illustrated in Fig. 3a, the summary food web can be reconstructed by first using prior knowledge to classify all 10 coexisting species as resource prey, meso predator, or top predator in a simple three-tier food web and then summarizing all potential interactions based on their trophic positions (see Methods). Following the procedure, a unique summary food web for the 10-species fish community in the LG pool was reconstructed and shown in Fig. 3b. In the food web, channel catfish and white bass are the top predators, freshwater drum and black crappie are the meso predators, and all other 6 fish species are resource prey. The summary network consists of 42 pairwise interactions (bidirectional links), among which 14 represent known predator-prey relationships (black arrows). Since the total possible number of pairwise interactions is 45 for 10 species, the summary food web does not impose sparsity on the interactions between fish species. These putative interactions can be naturally converted to the sign constraints of gLV model parameters (Fig. 3a, Additional file 2: Table S2): a positive, neutral, or negative interaction requires its corresponding parameter to be positive, 0 or negative as well.

**Fig. 3.**
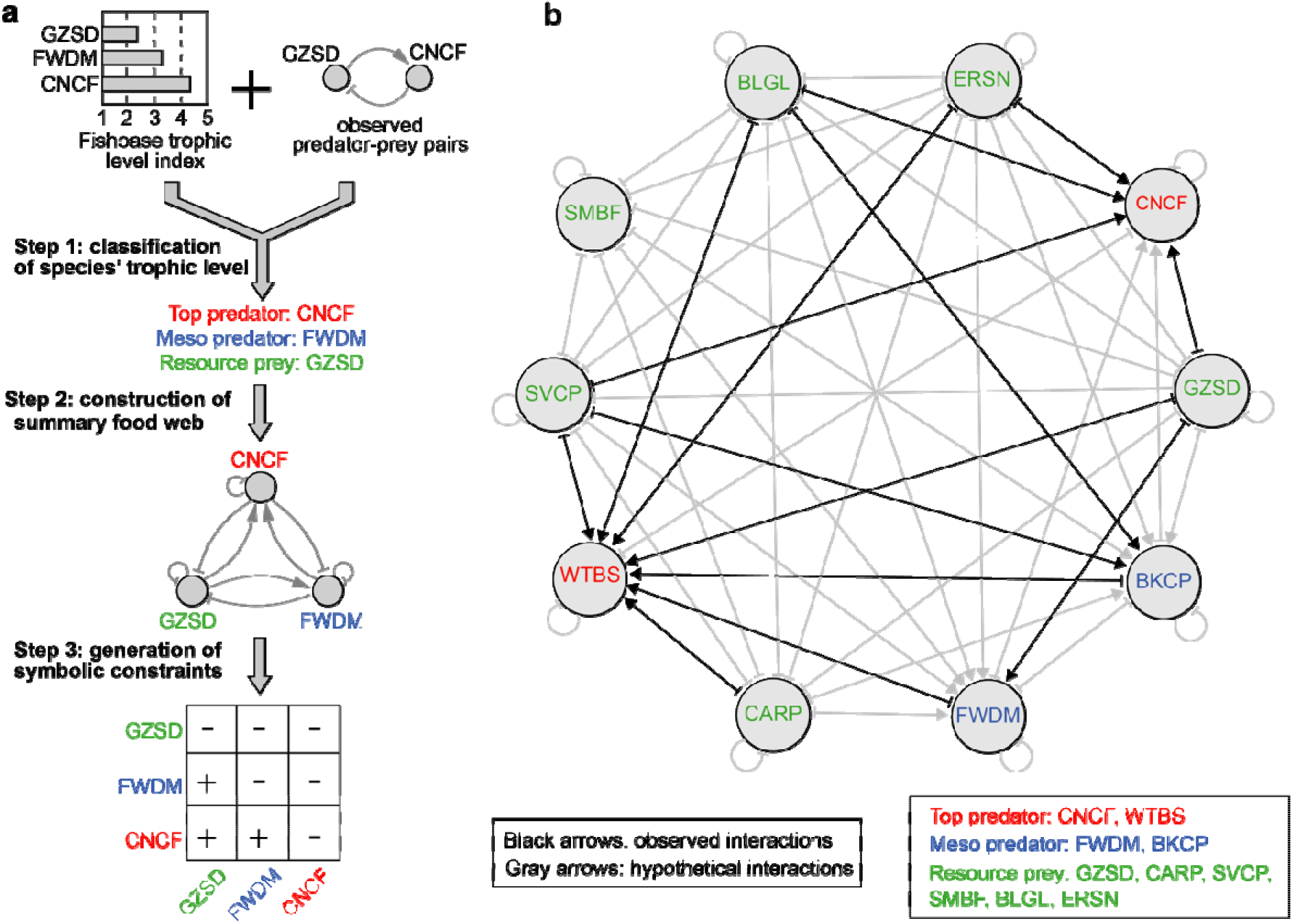
Construction of summary food web using parameter sign constraints. **a** Schematic illustration of a three-step procedure of generating symbolic constraints of interactions from prior knowledge (see Methods for details). **b** Reconstructed summary food web for the top 10 abundant fish species in the La Grange pool. Point arrows represent positive effects and blunt arrows represent negative effects. The observed predator-prey relationships in other water systems are indicated by black arrows, including BKCP-BLGL [42], CNCF-BLGL [43], CNCF-ERSN [31], CNCF-GZSD [31], FWDM-GZSD [44], WTBS-BKCP [45], WTBS-BLGL [5], WTBS-FWDM [45], WTBS-ERSN [46], WTBS-GZSD [46], WTBS-CARP [35] (the former species is a predator and the latter species is a prey).

### An ensemble of gLV models accounts for inference uncertainty

Our approach—which combines LGR with sign constraints—outperformed LR by improving adjusted R^2^ from 45% to 81% in fitting the fish abundance data from the LG pool (Additional file 1: Fig. S4). We excluded silver carp in the inference of growth rates and pairwise interaction coefficients for the 9 native species because the invasive species began to establish the Illinois River around 2000 [30] and has a much shorter time series. To prevent overfitting, we used empirical mode decomposition to smooth data (see Methods) and added a regularization term to the objective function (see Methods). An additional benefit of using smoothed data than original time-series is that LGR converged much faster (Additional file 1: Fig. S5).

If data are noise-free, the optimal fit should give the best estimate of network structure. However, uncertainty in data leads to uncertainty in parameter estimation so accounting for suboptimal yet constrained models can improve the inference power based on “the wisdom of crowds”. To search for alternative gLV models that are almost equally constrained by data, we generated a pool of 1000 perturbed models from the best-fit model given by LGR and constructed an ensemble by including only the subset with fitting error below a threshold (see Methods). Instead of using an arbitrary error cutoff, we found that the distribution of fitting errors of the 1000 models exhibited three well-separated peaks that naturally partition these models into three groups (Fig. 4a). Simulations of the 1000 models confirmed that their dynamics are very similar within the group (Fig. 4b) and the within-group mean adjusted R^2^ decreased from 79% for the first group to 61% and 2% for the second and third groups respectively. The superior performance of the first-group models simply assembled themselves into an ensemble that can be used for predictive analysis of fish community below.

**Fig. 4.**
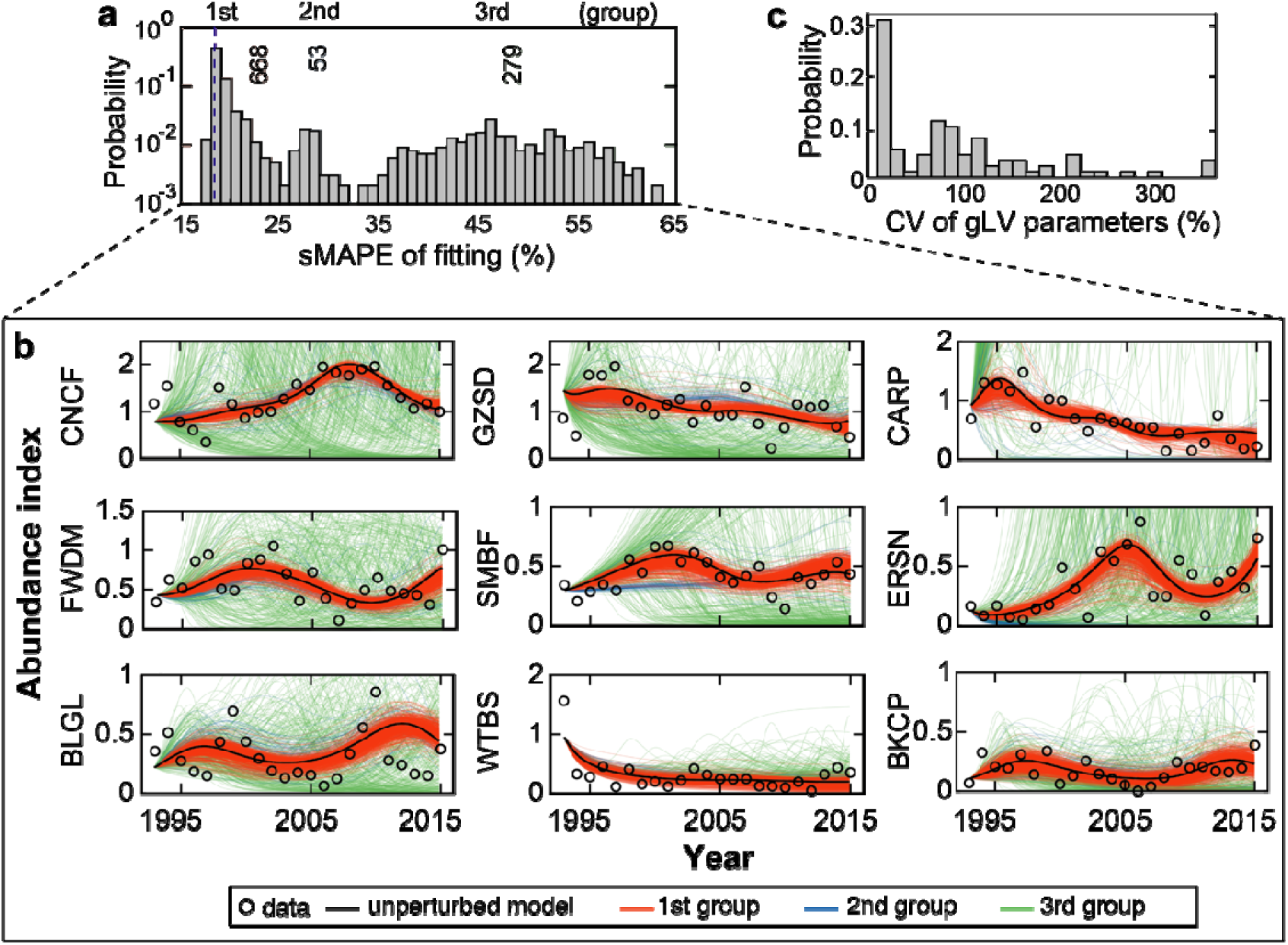
Ensemble method provides robust parameterization of generalized Lotka-Volterra (gLV) network models. **a** Probability distribution of the symmetric mean absolute percentage error (sMAPE) across 1000 gLV models perturbed from the best-fit model given by latent gradient regression (LGR). The distribution has three peaks that partition the 1000 models into three groups that represent good (668 models), mediocre (53 models) and poor (279) fits to data. Models in the first group were combined to make an ensemble. **b** Simulated trajectories of the fish abundance data by models from the three groups. Unperturbed model is the best-fit model given by LGR. **c** The coefficient of variation (CV) of gLV parameters across the 668 models in the ensemble.

### Probabilistic inference of native fish species’ growth and interactions

Using the ensemble, we quantified the extent of variability of gLV parameters (Additional file 2: Table S3) across its member models via the coefficient of variation (CV)—the standard deviation divided by the mean. The distribution of CV has a decreasing density (Fig. 4c) with 68% (36%) parameters of CV ≥ 0.25 (CV ≥ 1), suggesting large variability in the majority of parameters. Then we were wondering if their values inferred from data provide any evidence that the 9 native fish species grow and interact with each other. To answer this question, we tested the null hypothesis for each parameter of each individual ensemble member gLV model that its value is equal to zero. If the p-value of this test is *p*, then 1−*p* (what we call the “confidence score” below) informs how likely the parameter is different than 0 since its 100(1−*p*)% confidence interval just touches 0. In general, 1−*p* is proportional to the magnitude of its corresponding gLV parameter (Additional file 1: Fig. S6, Additional file 2: Table S4).

Averaging the confidence scores over the ensemble provides a more conservative measure of the evidences for species’ growth and interactions (Fig. 5a). The mean confidence scores for the per-capita growth rates of several prey (common carp, gizzard shad and emerald shiner) are 94%, 80% and 77% respectively, suggesting a high likelihood of their intrinsic population growth in the absence of other fish species. Although the mean confidence scores for almost all species interactions are low, the most probable interaction we inferred is a positive impact of emerald shiner on channel catfish with a 70% level of confidence, which agrees with empirical observations that emerald shiner support channel catfish’s growth by serving as major food sources [31]. To refine these predictions, we applied the same network inference procedure to fish abundance time-series data from the Pool 26—the closest pool to the LG pool (Fig. 1a) and had the most similar community composition (Fig. 1b). To include all 9 native fish species in the LG pool model, the pool 26 model must contain at least 12 species (Additional file 1: Fig. S2). We thus constructed an ensemble of 326 12-species gLV models (Additional file 1: Fig. S7, Additional file 2: Table S5, S6) with an ensemble-mean adjusted R^2^ 73%. The mean confidence scores estimated from the Pool 26 data identified with even higher possibility that emerald shiner grows in the absence of interactions (93%) and positively impacts channel catfish (72%) (Fig. 5b, Additional file 1: Fig. S7), thus confirming the predictions based on the LG data alone.

**Fig. 5.**
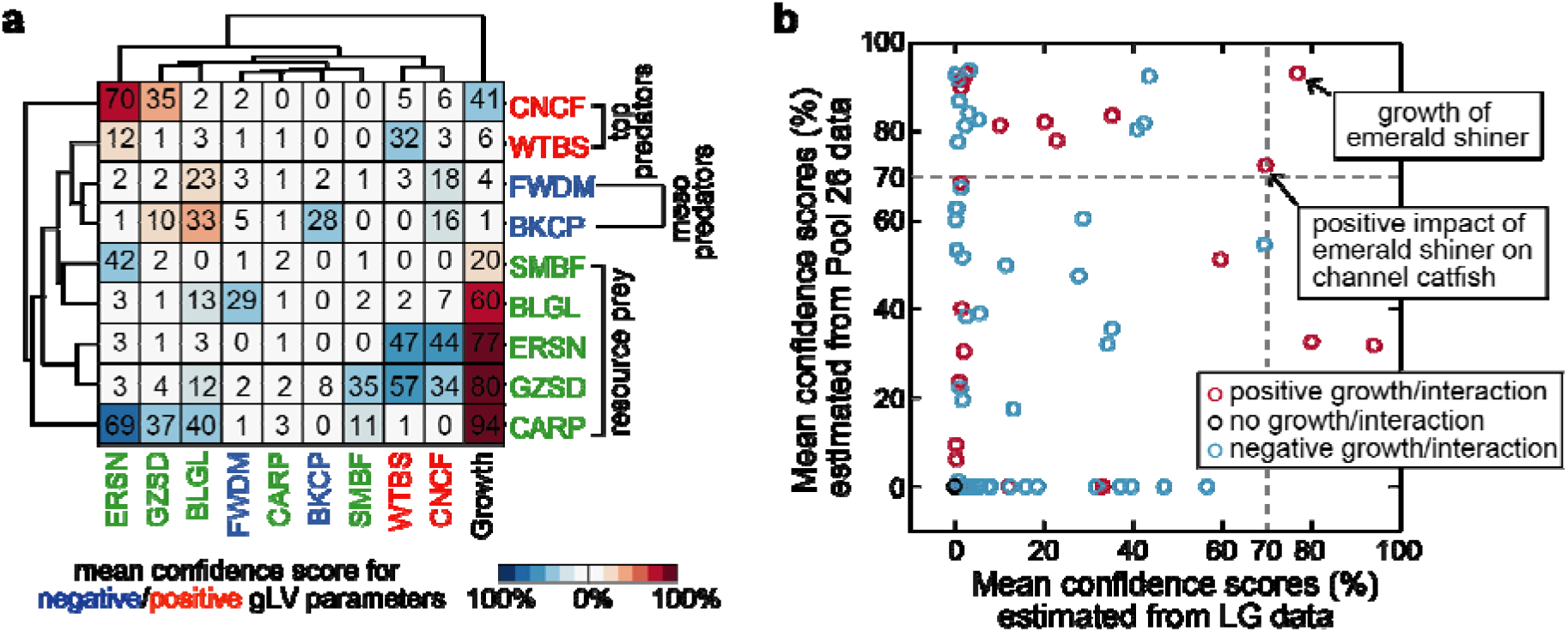
Mean confidence scores for species’ growth and interactions in the La Grange (LG) pool and the Pool 26. **a** Clustering of the mean confidence scores estimated from the LG data. The numbers in th square matrix made of the 9 rows and the first 9 columns are the mean confidence scores of pairwis interaction coefficients and indicate the likelihood that fish species on the column impacts fish species on the row. The numbers in the last column are the mean confidence scores of intrinsic growth rates and indicate the likelihood that population of each fish species grows (prey) or declines (predators) in th absence of the others. **b** Refinement of the predictions in **a** by combining mean confidence score estimated from both the LG and the Pool 26 data. Only the growth of emerald shiner and its positive impact on channel catfish have confidence scores at least 70% at both sites.

### Fluctuation of relative abundances of native fish species in the near future

Due to the decent accuracy of fitting existing data from the LG pool (adjusted R^2^ 79% on average), the ensemble of models was employed to predict the near future by extending their simulations for longer periods. In the next 20 years until 2035, the ensemble-mean trajectories of relative abundances show that 7 out of 9 dominant fish species in the LG pool fluctuate periodically and concordantly at the annual time scale (Fig. 6), suggesting that the LG pool fish community is a dynamically coupled ecosystem. In contrast, the relative abundances of the remaining two fish species, particularly the common carp, decreased continuously since the 1990s and were forecasted to remain at low level in the near future.

**Fig. 6.**
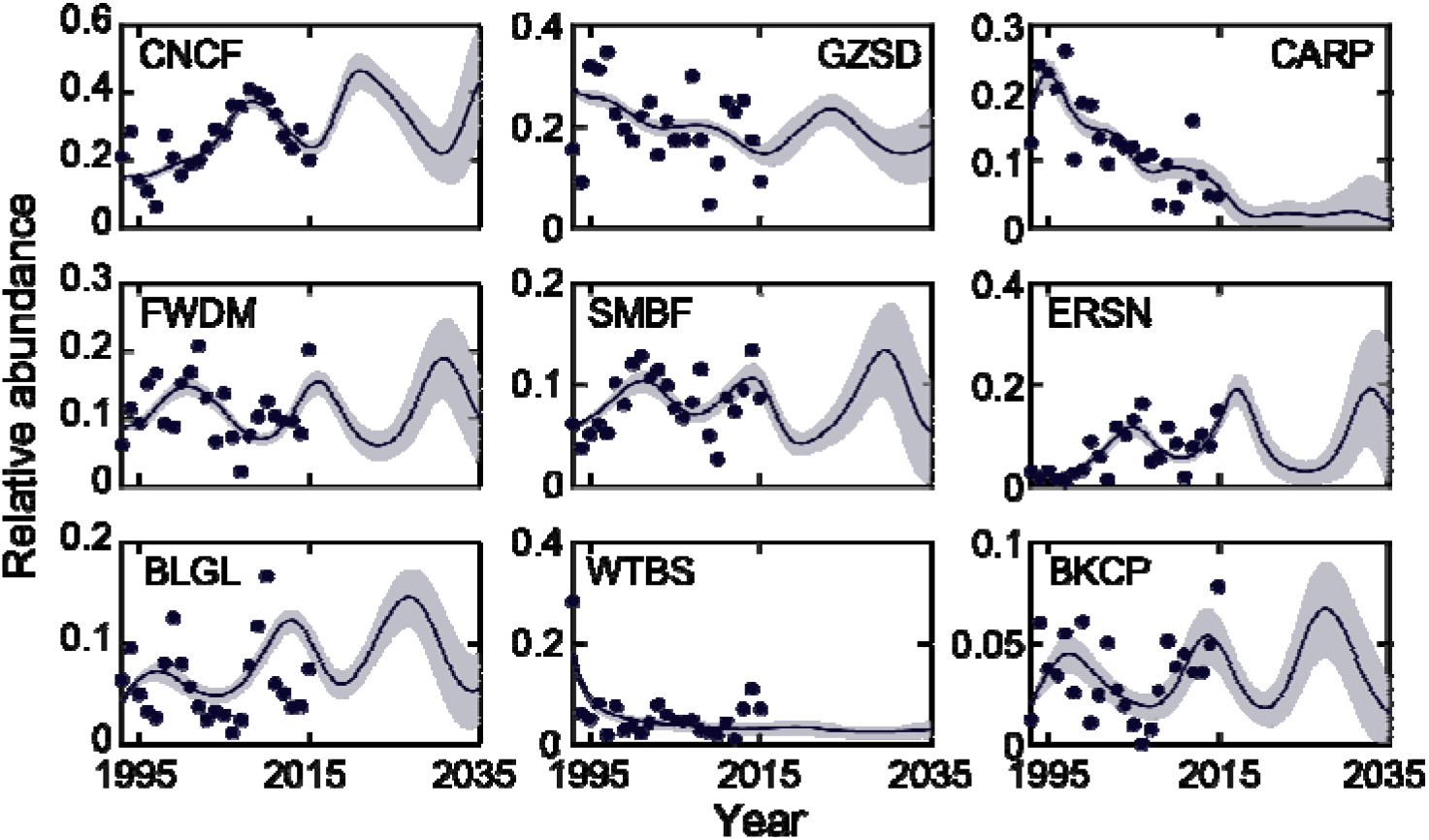
Forecasted population dynamics of the 9 dominant native fish species in the La Grange pool suggests a dynamically coupled ecosystem. Solid lines indicate the ensemble mean and gray shading indicate ensemble standard deviation. Filled circles: observed data.

### Impacts of invasive silver carp are stronger on native predators than prey

To study the impact of the silver carp—a present threat to the fisheries in the North America—we incorporated this species as a perturbation to the native fish network models in the LG pool. We assumed that its invasion altered the intrinsic growth rate of native fish species and quantified the susceptibility of each species to the perturbation using a single coefficient (see Methods). By fitting the susceptibility coefficients and testing whether their values are different than 0 for each gLV model in the ensemble (Additional file 2: Table S7, S8), we found stronger evidences that silver carp impacts native predators more than resource prey (Fig. 7). Particularly, the ensemble-mean confidence scores for the impacts of silver carp on the two top predators—channel catfish and white bass—are 78% and 91% respectively. Nonetheless, the confidences that the finesses of resource prey and even meso predators have been directly impacted by the silver carp are generally low, which justifies our earlier choice to exclude silver carp from the network inference.

**Fig. 7.**
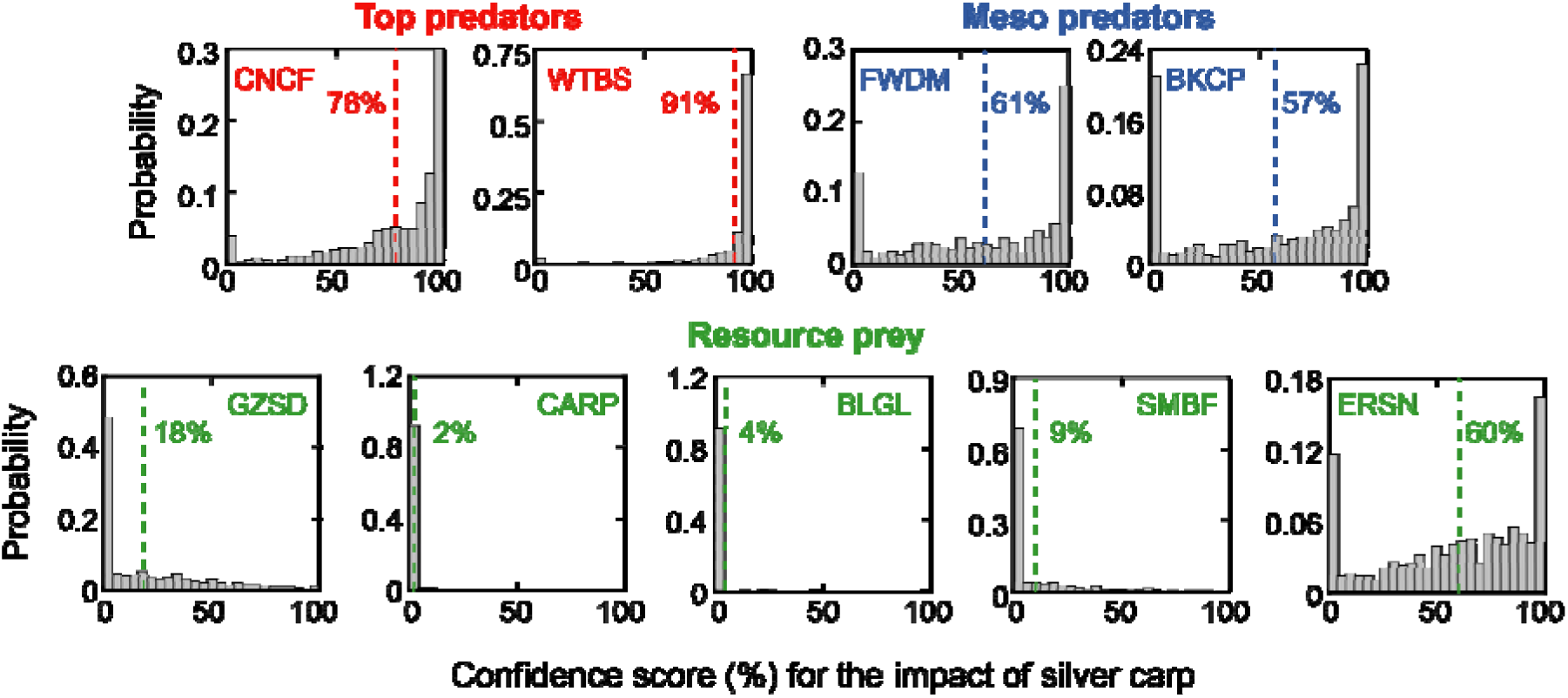
Probability distribution of the confidence scores for the impacts of silver carp on the 9 dominant native fish species in the La Grange pool. The scores associated with each native fish species indicate th likelihood that the impact from silver carp on this species is different than 0. The ensemble-mean of these scores are indicated by the dashed lines and the numbers beside them.

## Discussion

Here we proposed a new method to infer ecological networks from field data on real-world ecosystems. Field data are invaluable for ecology, but noise and infrequent sampling hinders network inference—especially with population dynamics models such as gLV which requires the calculation of time gradients [11]. The problem could in principle be solved by measuring accurate data and at higher rates, but this is often impractical. The inference method we proposed here offers a practical solution based on a deterministic optimization algorithm combined with parameter sign constraints obtained from prior knowledge and an ensemble method to assess the uncertainty associated with deterministic predictions. Modeling time gradients as latent parameters could improve other inference algorithms, especially those mathematically equivalent to gLV such as the Ecopath modeling framework [32].

It is interesting to observe from data that the relative abundance of common carp has decreased over time since the 1990s (Fig. 6). First introduced to the United States since 1800s, common carp were initially more competitive than native competitors because they reproduced rapidly and can survive in poor water quality [33]. Since its intrinsic growth rate is very likely to be positive (94% confident; see Fig. 5a), the declined relative abundance of common carp may be due to stronger competitive inhibitions from native consumers in the past several decades. Particularly, a moderate-level evidence (69%) was assigned to the inhibition of common carp by emerald shiner (Fig. 5a). Emerald shiner is a small fish species feeding on a variety of zooplankton, protozoans and diatoms. Considering its growth and impact on channel catfish were the only gLV coefficients identified with ≥70% confidence at both the LG pool and the Pool 26, emerald shiner might be a keystone species that drives changes in the relative abundance of local fish communities.

Our results also suggested that the ecological consequences caused by the silver carp’s invasion may not be too detrimental in short-term period. Overall, we found little evidences that the invasion had impacted the fitness of native prey fish. The lack of strong negative impacts of silver carp on native resource prey may be due to the high productivity and species richness in the Illinois River [34], which mitigates the effects of interspecific competition for food sources. Still, we estimated, with 78% and 91% confidences respectively, that channel catfish and white bass may eat silver carp and benefit from supplemental prey that they catch. These findings are consistent with stomach content analysis of native predators in the LG pool—including channel catfish, black crappie, and white bass, which revealed that silver carp had indeed entered their diets by serving as alternative prey [35].

Our study has limitations that stem from both the limitations of gLV model and the inference approach we developed. The gLV model has known limitations, including additivity (fitness influence that each species receives from others is additive) and universality (the sign and strength of the influence can be reflected by the interaction coefficient) assumptions [36], linear functional responses (efficiency of predation is unsaturated even when the prey is very abundant) [37], and the paradigm of pairwise interactions between species (high-order interactions are not considered) [38]. These limitations can be in principle overcome by increasing model complexity such as using saturated functional responses, which would nonetheless abolish the benefits associated with linear transformation of gLV equations during parameterization.

Our inference method has additional limitations. First, the major predictions made using a criterion of “70% confidence at both sites of the LG pool and Pool 26” may lead to type I errors. However, this is expected given insufficient and noisy data. Second, the LGR algorithm is a local optimization approach that easily falls into local minima; there is no guarantee that the optimized gLV parameters are closer to the ground truth (if it exists) than the initial guesses. This limitation has been reflected in our benchmark test where parameters that fit the data better could be further from the truth (Fig. 2c). Since the output of LGR depends on initial guesses which further depend on data, the issue of local optimization can also lead to instability of the algorithm in cross validation with random partitioning of the data into the training and testing subsets. Although global optimization techniques such as Markov chain Monte Carlo may diminish the limitation, they generally require intensive computations. Third, LGR may fail numerically in the step of solving a gLV model when its parameters are not well constrained and cause the simulation to explode. Therefore, a robust use of LGR requires parameter constraints such as the sign constraints we derived from a summary food web (Fig. 3b). However, this is only one way to incorporate prior knowledge and other types of constraints may be imposed to reduce the number of interactions further. Lastly, environmental factors such as temperature were not considered but they can be easily added as exogenous variables (similar to the silver carp) in the future.

## Conclusions

We advanced the gLV model-based network inference and showed its utility in inferring/predicting the network structure and dynamics of a freshwater fish community in the Illinois River. Future applications of the inference approach could be generalized to study fish communities in other geographical locations with varying ecological and environmental conditions (e.g., other rivers with long-term resource monitoring data) or even other macroscopic organisms. Such applications may enhance the ability to understand and predict the structure and dynamics of natural ecosystems and shed light on disruptive threats posed by invasive species.

## Methods

### General

All simulations and computational analyses were performed in MATLAB R2018 (The MathWorks, Inc., Natick, MA, USA).

### Long term resource monitoring data

The time series data of the Upper Mississippi and Illinois Rivers fish community were collected from the annual reports of the Long Term Resource Monitoring Program [22]. The program used a multigear and multihabitat sampling design protocol (refer to the program report for details) to collect data from 6 observation sites (Lake City, Minnesota, Pool 4; La Crosse, Wisconsin, Pool 8; Bellevue, Iowa, Pool 13; Alton, Illinois, Pool 26; Havana, Illinois, La Grange Pool; and Cape Girardeau, Missouri, Open River). To standardize the catch per unit effort (CPUE) from multiple gears to the same relative scale, the raw CPUE data during the time period between 1993 and 2015 were converted to relative abundance among species within the same site and summed over all 6 fishing gears (electrofishing, fyke net, mini fyke net, large hoop net, small hoop net, trawling). Since the absolute abundances are not available, we assumed that the fish species were maintained at or nearby the carrying capacity, which allows for parameterizing a generalized Lotka-Volterra model directly from relative abundance data such as the standardized CPUE indices.

### Noise filtering and data smoothing

It is well known that outliers or noisy data in the population abundance data can result in spurious gradient estimates. Although our parameter estimation algorithm was designed to solve this issue by optimizing the gradients, it is nonetheless a local optimization approach and uses the numerically approximated gradients as initial guesses to start the fitting procedure. To improve the fitting robustness, population abundance data for the two microbial ecosystems as well as the two fish communities in the La Grange pool and the Pool 26 were smoothed before used to guide parameterization.

Data smoothing was performed by the classical empirical mode decomposition (EMD) algorithm which has been extensively reviewed elsewhere [39]. Briefly, EMD decomposes the input time-series data into several intrinsic mode functions (IMF), each of which represents a distinct local oscillation mode of the data. Since IMFs with Hurst exponent below 0.5 have low autocorrelations and are more likely to contain noise than signal, smooth trends can be extracted from the original time-series by only keeping IMFs with Hurst exponent no smaller than 0.5. The MATLAB codes of EMD and Hurst exponent estimation can be accessed from https://www.mathworks.com/matlabcentral/fileexchange/52502-denoising-signals-using-empirical-mode-decomposition-and-hurst-analysis.

### Generalized Lotka-Volterra model

The generalized Lotka-Volterra (gLV) model is a system of ordinary differential equations (ODE) with birth-death processes describing how fish species abundances change over time

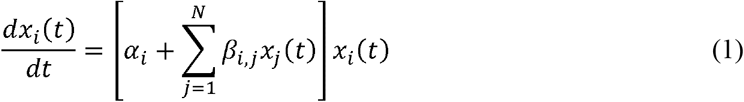

where *x*_*i*_(*t*) is the abundance of fish species *i* at time *t* and *N* is the total number of fish species. *α*_*i*_ is referred to as the net (birth minus death) population’s per-capita growth rate of the fish species *i* while *β*_*i,j*_, known as the pairwise interaction coefficient, represents the population influence of fish species *j* on fish species *i*. Once parameterized, Equation (1) can be numerically solved using any ODE solver. We used MATLAB’s built-in solver *ode15s* in this study.

### GLV parameterization by linear regression (LR)

A commonly used technique to parameterize a gLV model is to discretize Equation (1) and solve the following multilinear regression [11]

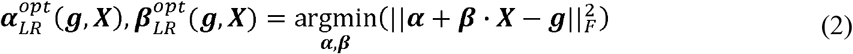

where ‖ · ‖_*F*_ is the Frobenius norm. ***α***, ***β***, ***X***, ***g*** are the vectors/matrices of growth rates, interaction coefficients, time-series data, and gradients of the time-series data respectively (*t*_1_, *t*_2_,…,*t*_*M*_ are discrete time points)

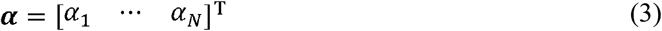

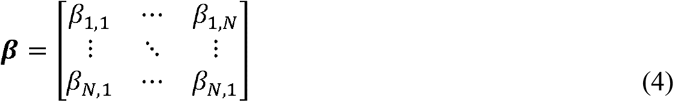

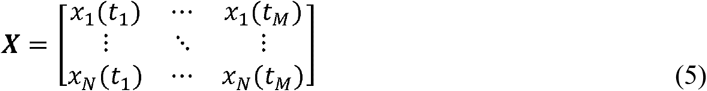

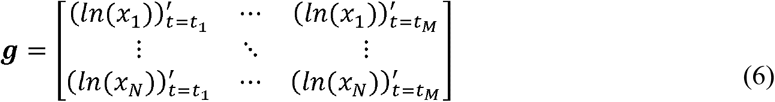

Note that the gradients ***g*** are input parameters to the linear regression procedure and need to be numerically approximated. We calculated ***g*** by differentiating the spline interpolants of the observed data ***X***. MATLAB built-in function *spline* and *fnder* were used for spline interpolation and differentiation respectively. The linear least-square problem in Equation (2) was solved by the interior-point algorithm implemented by MATLAB built-in function *lsqlin*.

### GLV parameterization by nonlinear regression (NLR)

The gLV parameters ***α***, ***β*** can also be estimated by nonlinear regression. Naively, it searches the space of ***α***, ***β*** for a local minimum of a sum of squares between simulated and observed data

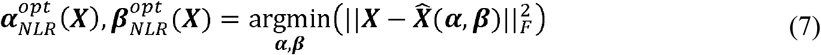

where 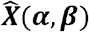 is the matrix that has the same format as ***X*** but consists of simulated time-series data 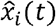 obtained by numerically solving the gLV model with given ***α***, ***β***, i.e.,

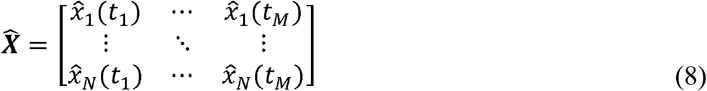

The nonlinear least-square problem in Equation (7) was solved using the trust-region-reflective algorithm, which was implemented by MATLAB built-in function *lsqnonlin*.

### GLV parameterization by latent gradient regression (LGR)

Our approach minimizes the same least square as in NLR but searches the space of the latent gradients ***g***, rather than gLV parameters ***α***, ***β***

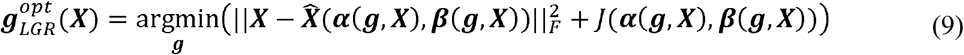

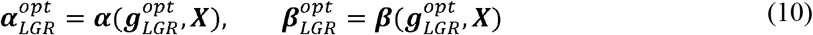

The transformation functions ***α***(***g***, ***X***), ***β***,(***g***, ***X***) can be found by solving the linear regression in Equation (2), i.e., 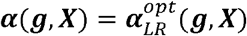 and 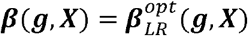. *J*(***α***, ***β***) in Equation (9) was introduced as the penalty function to reduce the risk of overfitting. Here we used a modified version of ridge regression where the self-interaction coefficients of species are not penalized (this is consistent with our previous assumption that the fish community is saturated nearby carrying capacity, which implies strong intraspecific competitions)

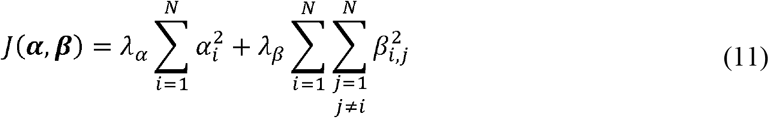

where λ_*α*_ and λ_*β*_ are the penalty coefficients for the growth rate vectors and the interaction matrix respectively.

The number of observed data is much larger than the number of parameters for synthetic ecosystem and the two microbial ecosystems. Therefore, we used λ_*α*_ = λ_*β*_ = 0 in fitting these data. For the fish abundance data in the LG pool and the Pool 26, we performed leave-one-out cross-validation: the training dataset was the full time-series excluding the middle-year data (*t*_*test*_ = 2004) and the test dataset includes a single data point at that year. As we mentioned in the Discussion section, both local optimization nature of LGR and insufficient data prevented us from using more complex strategies of data partitioning between training and testing sets. The optimal values of λ_*α*_ and λ_*β*_ were chosen as the combination minimizing the sum of squared error over all fish species on the test set, i.e., 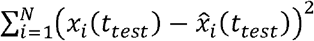. We found λ_*α*_ = 1.6 × 10^−4^, λ_*β*_ = 7.9 × 10^−3^ for the LG pool data and λ_*α*_ = 1.6 × 10^−2^, λ_*β*_ = 4.0 × 10^−4^ for the Pool 26 data. The final gLV model was parameterized by running LGR with the optimized penalty coefficients and the complete dataset.

Solving Equation (9) requires an iteration method that alternates between updating the values of ***g*** and ***α***, ***β***. The algorithm of LGR include 4 distinct steps

1. Pick an initial guess of ***g***^(0)^ for ***g***. We constructed ***g***^(0)^ by numerical differentiation of data as described above (see **GLV parameterization by linear regression** for details).
2. Given ***g***^(*k*−1)^ and ***X***, estimate ***α***^(*k*)^, ***β***^(*k*)^ by solving the following linear regression

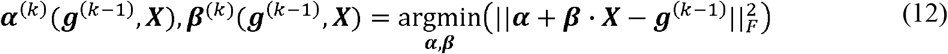
3. Given ***g***^(*k*−1)^, ***α***^(*k*)^, ***β***^(*k*)^ and ***X***, estimate ***g***^(*k*)^ by applying the update rule of the Levenberg-Marquardt (LM) algorithm [40] (other optimization algorithms can be applied similarly). Let ***X***_1_, 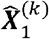, 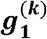 are the flattened 1-dimensional *NM* × 1 vectors of ***X***, 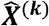, and ***g***^(*k*)^ respectively. The LM algorithm is a blend of the gradient descent and a Gauss-Newton approach that constructs a search direction by solving the following set of linear equations

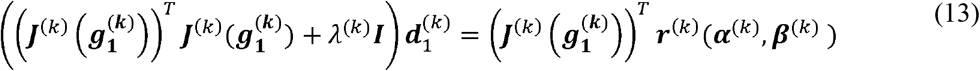

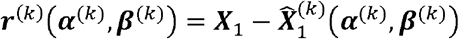 is the *NM* × 1 residual between observed and simulated data. 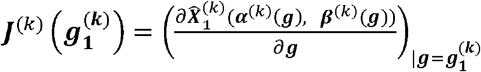 is the *NM* × *NM* Jacobian matrix. λ^(*k*)^ is a damping parameter that controls the magnitude and direction of the update (small values of λ^(*k*)^ result in a Gauss-Newton update and large values of λ^(*k*)^ result in a gradient descent update). ***I*** is the identify matrix. Let ***d***^(*k*)^ be the reshaped 2-dimensioanl *N* × *M* matrix of 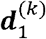. The update rule of the LM algorithm can be represented as below

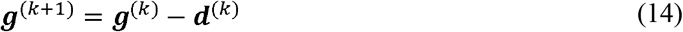
4. Let *k* = *k* + 1 and go back to step 2. The iterations continue until the convergence criteria for the LM algorithm is met.

The LM algorithm is implemented by MATLAB built-in function *lsqnonlin*. The choice of λ^(*k*)^ at each step and more details about the implementation are available in the MATLAB webpage https://www.mathworks.com/help/optim/ug/least-squares-model-fitting-algorithms.html#f204.

The above iterative optimization procedure is a deterministic variant of the expectation-maximization algorithm [41]. The latent gradients computed in the expectation step (Step 3) are used to update the gLV coefficients in the maximization step (Step 2). However, our approach was not formulated into a statistical framework that explicitly models the gLV parameters and the latent gradients as random variables with probabilistic distributions. Therefore, it is still a deterministic optimization method that should not be confused with a classical expectation-maximization algorithm.

### Synthetic community data

To benchmark our LGR algorithm, we created a 3-species (*X*_*i*_ where *i* = 1,2,3) gLV model with its parameter values (*α*_*i*_ and *β*_*i,j*_ where *i,j*= 1,2,3) indicated along the arrows in the model diagram (Fig. 2b). The synthetic data used in Fig. 2b were created by deterministically solving the model using MATLAB built-in function *ode15s*. Environmental noise was added to the model by simulating stochastic differential equations

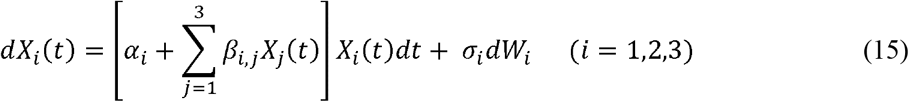

where *dt* is the time step and *dW*_*i*_ is the Wiener process (Brownian motion) with diffusion rate *σ*_*i*_ (equal to 0.001 for all three species). The histograms in Fig. 2c were plotted based on 1000 simulated noisy datasets. The MATLAB codes for numerical solution of stochastic differential equations can be assessed from https://github.com/horchler/SDETools.

The following setups are general to both deterministic and stochastic simulations. First, synthetic data used in Fig. 2b,c and Additional file 1: Fig. S3 were generated by sampling the simulated trajectories at a fixed time interval of 5 from *t* = 0 to *t* = 100. Second, the initial conditions for *X*_1_, *X*_2_, *X*_3_ in all simulations were 0.15, 0.6 and 0.4 respectively. Lastly, parameter sign constraints were employed by all inference algorithms (LR, NLR, LGR) in fitting the synthetic data.

### Summary food web and parameter sign constraints

The summary food web of the modelled fish community was reconstructed in two steps: (1) classifying all fish species in three trophic levels represented by resource prey, meso predator, and top predator on the basis of their feeding behavior; (2) summarizing all potential interactions based on the classification and empirical observations. In the classification step, the trophic positions of fish species was determined by finding a distribution that is compatible with two constraints imposed by prior data: (1) the FishBase (https://www.fishbase.de) trophic level index (a floating-point number that equals one plus weighted mean trophic level index of the food items) of any fish species in higher trophic levels is no smaller than that of any fish species in lower levels; (2) the predator of any known predator-prey relationship occupies a higher trophic level than the level occupied by the prey. We assume that each pair observed to interact in other freshwater ecosystems has the potential to interact the same way in the Upper Mississippi and Illinois Rivers.

In the summarization step, the potential pairwise interactions include not only observed predator-prey relationships but hypothetical interactions that are generated by the following ecological rules: (1) fish species on higher trophic levels feed on fish species on the immediate lower level (common prey relationships); (2) the same fish species compete for limited resources within its own population (intraspecific competitions); (3) fish species on the same trophic level compete with each other for limited resources (interspecific competitions). Any pair of fish species whose trophic relationship does not apply to the three rules is assumed to be non-interacting.

Sign constraints can be converted from the potential interactions in the summary food web. Depending on the interaction type, the conversion follows the following rules: (1) *β*_*i,j*_ < 0 and *β*_*j,i*_ > 0 for predator (species *j*)-prey (species *i*) relationships; (2) *β*_*i,i*_ < 0 for intraspecific competitions within population of species *i*; (3) *β*_*i,j*_ < 0 and *β*_*j,i*_ < 0 for interspecific competitions between species *j* and species *i*; (4) *β*_*i,j*_ = 0 and *β*_*j,i*_ = 0 for non-interacting species pairs. Per-capita growth rate of species *i* is positive (*α*_*i*_ > 0) if it occupies the lowest trophic level and negative (*α*_*i*_ < 0) if it occupies higher trophic levels. The derived sign constraints for the La Grange pool and the Pool 26 were combined and shown in Additional file 2: Table S2.

### Construction of ensemble models

To identify alternative parameters that fit data (almost) equally well, we first generated perturbed gLV coefficients by adding noise to the coefficients 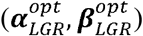 of the optimal (unperturbed) model obtained by LGR. Noise was added by sampling a log-normal distribution with the mean equal to the logarithmic of 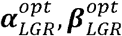 and the standard deviation fixed at a constant *σ*. Then the perturbed coefficients were used as initial guesses and re-optimized to minimize the following regularized least-square objective function

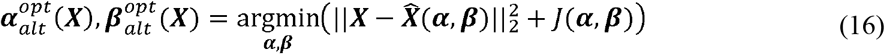

where 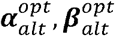 are gLV coefficients of the re-optimized model. The MATLAB trust-region-reflective algorithm was used to solve the above nonlinear regression. The standard deviation (*σ*) of the lognormal distribution was carefully chosen to ensure that the deviations of the re-optimized models from the data span a distribution that is neither too wide (low sampling efficiency) nor too narrow (not enough diversity). We found that *σ* = 0.2 and *σ* = 0.005 serve the purpose for the LG pool and the Pool 26 respectively.

For each of the LG pool and the Pool 26, we generated 1000 perturbed-then-reoptimized models as candidates for building an ensemble of models that fit data (almost) equally well. Practically, we used a cutoff value to exclude those models whose deviations from the data are higher than a threshold. In Fig. 4a, we quantified the deviation of model from data using symmetric mean absolute percentage error (sMAPE)

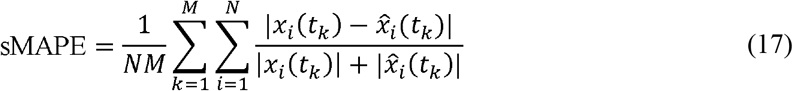

where *x*_*i*_(*t*_*k*_) and 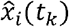 are observed and simulated abundance of fish species *i* at time *t*_*k*_. We preferred sMAPE over other metrics such as the mean squared error because (1) it is normalized between 0 and 1 and (2) more importantly, its distribution over the 1000 models for the LG fish community provides a less arbitrary cutoff value (0.25) that separates candidate models into groups that represent good and poor fits to data (Fig. 4a). To ensure fair comparison between model predictions across observation sites, we applied the same cutoff criterion (sMAPE ≤ 0.25) to construct the ensemble of gLV models for the Pool 26 fish community.

### Silver carp models

We chose not to model the abundance of silver carp as an autonomous gLV variable because the number of data points in silver carp’s time series was insufficient to reliably estimate new gLV parameters associated with the variable. Instead, silver carp was added to the gLV models of the native fish network in the LG pool as an exogenous variable (its value is not determined by model but given by data)

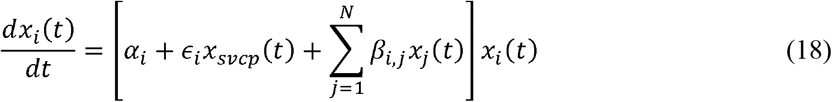

where *ϵ*_*i*_ is the susceptibility parameter that quantifies the response of the growth of native fish species *i* to silver carp. *x*_*svcp*_(*t*) is the abundance of silver carp at any time *t*, which can be obtained by interpolating data observed at discrete time points. Since silver carp invaded the Illinois River for only two decades, we assumed that silver carp perturbs the growth rate of native fish species without changing their feeding behavior and interactions with other native species. In other words, the coefficients *α*_*i*_ and *β*_*i,j*_ inferred in the absence of silver carp remain unchanged in its presence. For each ensemble gLV model with parameters 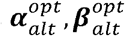, the optimal value of its susceptibility parameter 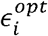 was given by the following nonlinear regression

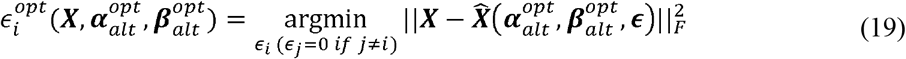

where ***ϵ*** = [*ϵ*_1_ ··· *ϵ*_*N*_]^T^. Note that we fit each *ϵ*_*i*_ one at a time while setting all other *ϵ*_*j* (*j*≠*i*)_ to zero, since too many adjustable parameters may lead to overfitting and spurious coupling. Equation (19) was solved using trust-region-reflective algorithm implemented in *lsqnonlin*, together with sign constraints of ***ϵ*** (Fig. 3b, Additional file 2: Table S2).

### Confidence score

The confidence score of a parameter is defined as 1 minus p-value testing that the parameter value is different than 0, i.e., 1 minus the minimum significance level below which the confidence interval of the parameter includes 0. If ***z*** is the vector of parameters (it could be gLV parameters ***α***, ***β*** in Equation (1) or susceptibility parameters ***ϵ*** in Equation (18)), its confidence interval at significance level *α* is given by

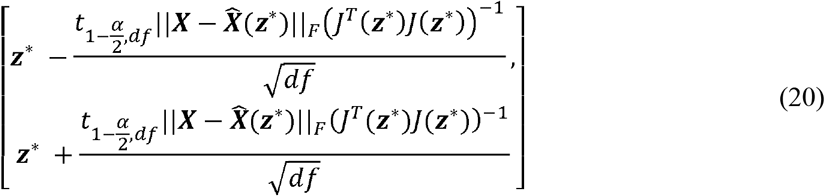

***z***\* is the optimized value of ***z***, *df* is degree of freedom (number of data minus number of parameters), ***X*** and 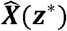 are the observed and simulated data respectively, 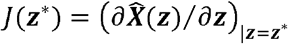 is the Jacobian evaluated at ***z***\*, and 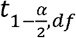 is the Student’s *t*, inverse cumulative distribution function. We used MATLAB built-in function *nlparci* to construct confidence intervals (*nlparci* essentially computes Equation (20)).

## Declarations

## Abbreviations

gLV: generalized Lotka-Volterra
LR: linear regression
NLR: nonlinear regression
LGR: latent gradient regression
LM: Levenberg-Marquardt
CV: coefficient of variation
sMAPE: symmetric mean absolute percentage error
MSE: mean squared error
LG: La Grange
OR: Open River
CPUE: catch per unit effort
EMD: empirical mode decomposition
IMF: intrinsic model function
PCA: principle component analysis
CNCF: channel catfish
GZSD: gizzard shad
CARP: common capr
FWDM: freshwater drum
SMBF: smallmouth buffalo
ERSN: emerald shiner
BLGL: bluegill
WTBS: white bass
BKCP: black crappie
SVCP: silver carp

## Ethics approval and consent to participate

Not applicable.

## Consent for publication

Not applicable.

## Availability of data and materials

The raw fish abundance data in all six observation sites can be accessed from the website of the Upper Mississippi River Restoration Program (https://umesc.usgs.gov/field_stations/fieldstations.html). The standardized CPUE indices for the six sites are available in Additional file 3. The MATLAB scripts for latent gradient regression have been submitted to https://github.com/liaochen1988/fish_network_inference. Other data supporting the findings of this study are available from either Additional files or the corresponding author on reasonable request.

## Authors’ contributions

C.L and Z.Z. conceived the study. C.L. developed the inference method and implemented simulations; C.L and Z.Z. wrote the initial draft; J.X. reviewed and edited subsequent versions of the manuscript; All authors designed the research, discussed the results and analyzed data.

## Acknowledgements

Not applicable.

## Additional files

Additional file 1.docx: Supplementary Figures.

Additional file 2.xlsx: Supplementary Tables.

Additional file 3.xlsx: Standardized fish abundance indices for all six observation sites in the Upper Mississippi and Illinois Rivers.

