## Additional file 1 for "Enhanced inference of ecological networks by parameterizing ensembles of population dynamics models constrained with prior knowledge"

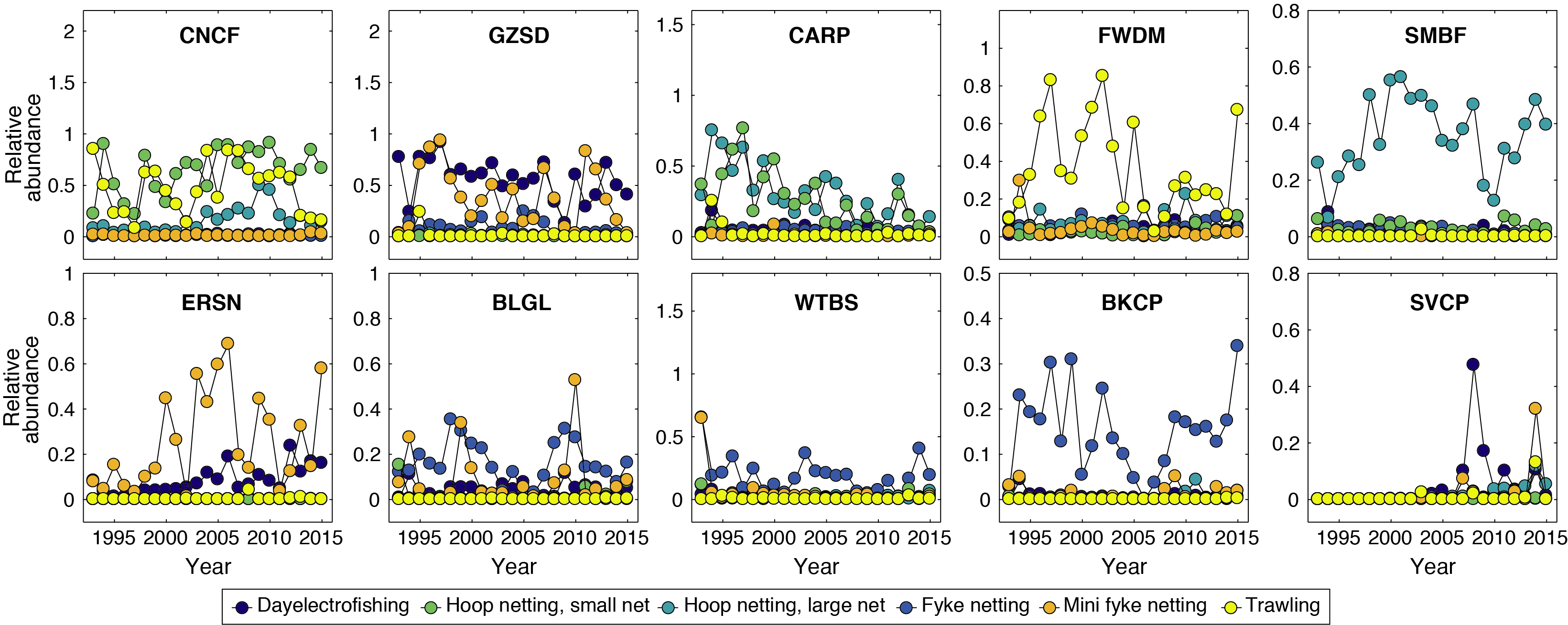

**Fig. S1** Relative abundance (normalized among species) from multiple fishing gears for the dominant fish species in the La Grange pool. Different fishing gears are selective for certain types of fish and no gear can catch all of them, suggesting the need to combine fish samples from different gears. Therefore, these relative abundances were summed over fishing gears to obtain standardized CPUE (catch per unit effort) indices which were later used to fit generalized Lotka-Volterra model.

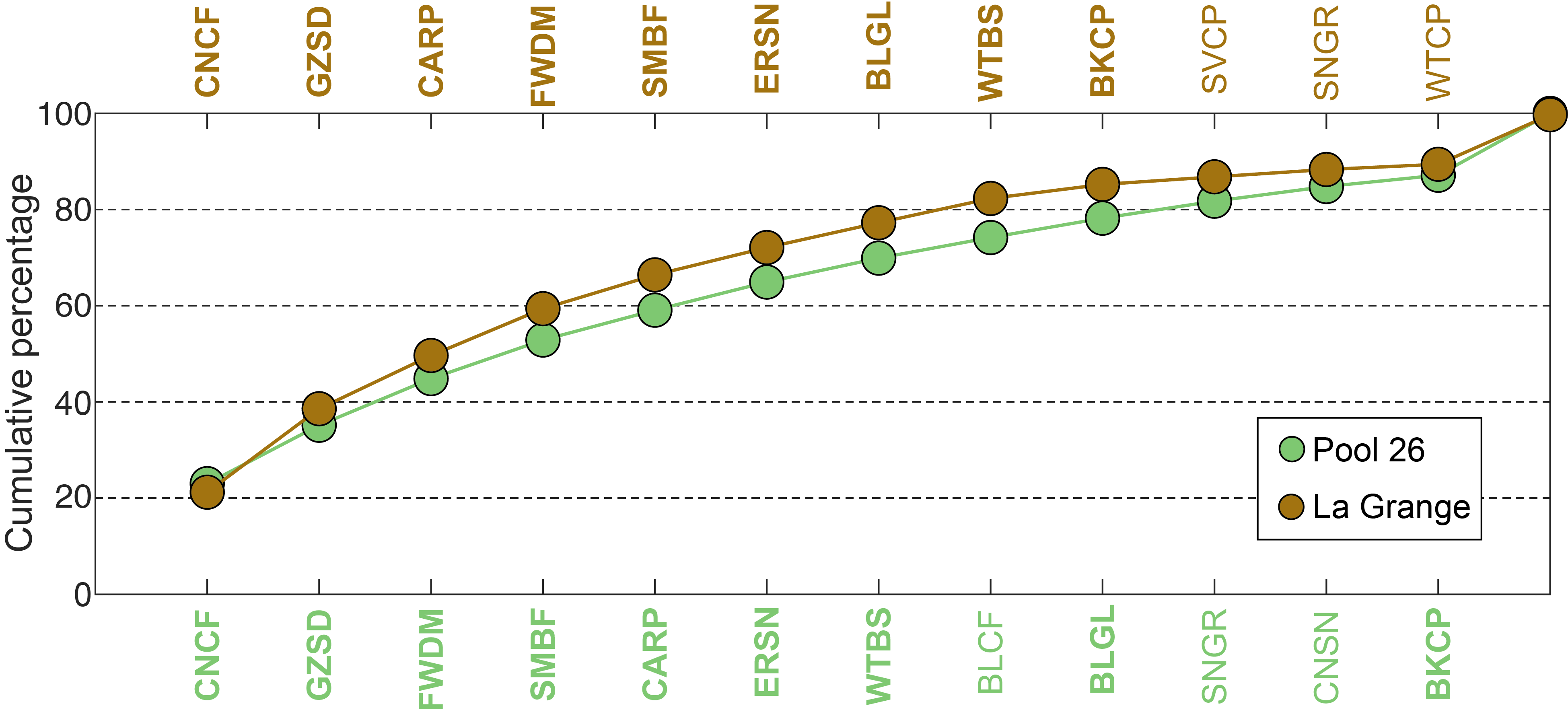

**Fig. S2** Cumulative percentage of averaged (between 1993 and 2015) abundance index. The top 12 dominant fish species at both the La Grange pool and the Pool 26 are shown in the order of most to least abundant, where fish species common to both sites are made bold.

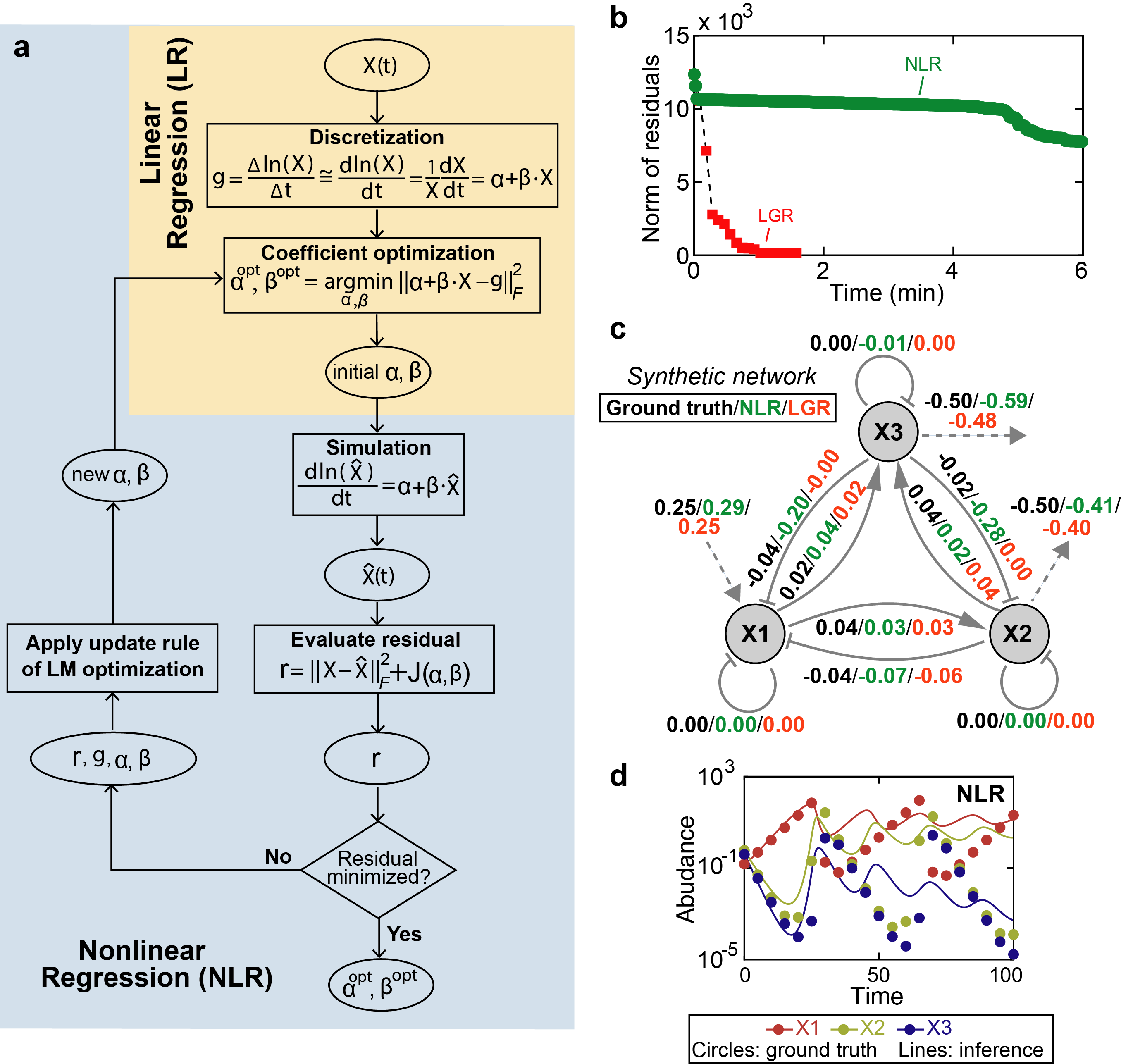

**Fig. S3** Comparison of nonlinear regression (NLR) and latent gradient regression (LGR) algorithms for fitting synthetic data generated by the same model as in the main text Fig. 2b. **a** Flow chart of nonlinear regression. $X\left( t \right)$: observed time series; $\hat{X}\left( t \right)$: simulated time series; $\alpha,\beta$: parameters of the generalized Lotka-Volterra (gLV) model; $g$: gradients (i.e., time-derivatives of $\ln(X\left( t \right))$; $J(\alpha,\beta)$: penalty function; $||\cdot||_{F}$: Frobenius norm; LM: Levenberg-Marquardt. Nonlinear optimization takes the output of linear regression as initial guesses of $\alpha,\beta$ and adaptively update $\alpha,\beta$ until the difference between observed and simulated time series is minimized. **b-d** Comparison between NLR and LGR for their convergence rates (**b**), inferred parameter values (**c**) and fitted trajectories (**d**). Each symbol (diamond and cross) in **b** marks the time of completion of one optimization iteration. In **c**, solid arrows represent positive (point end)/negative (blunt end) interactions and dashed arrows represent intrinsic population growth (incoming)/decline (outgoing) in the absence of other species. Ground-truth and inferred values of both interaction strengths and per-capita growth rates are indicated along the network links.

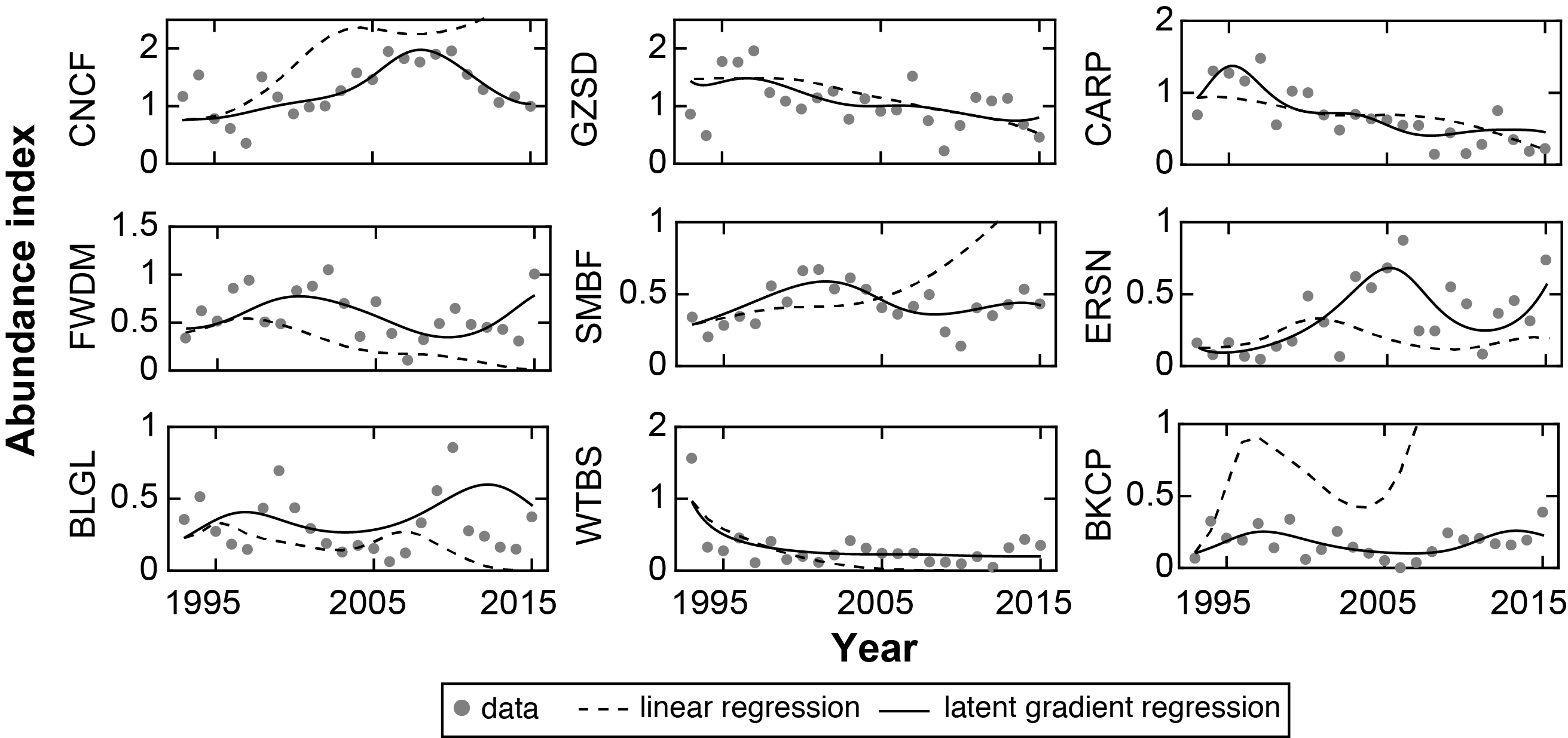

**Fig. S4** Comparison of linear regression and latent gradient regression algorithm in fitting fish abundance data for the 9 dominant fish species in the La Grange pool. Solid and broken lines: simulation; gray dots: observed data.

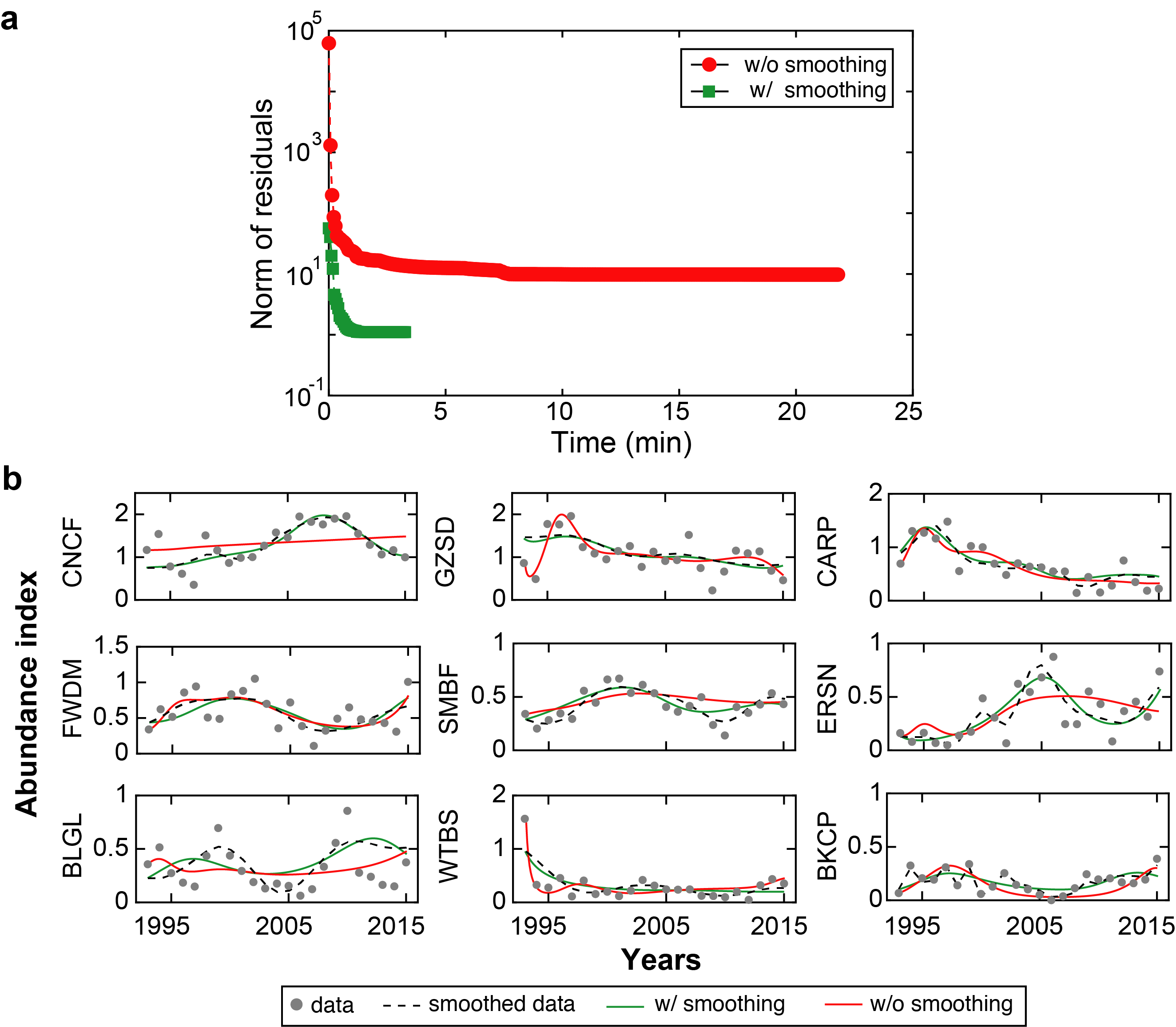

**Fig. S5** Effects of smoothing on the convergence speed (**a**) and accuracy (**b**) of fitting abundance data for the La Grange pool fish community using latent gradient regression. The adjusted R^2^ of fitting is the same (81%) between w/ and w/o smoothing.

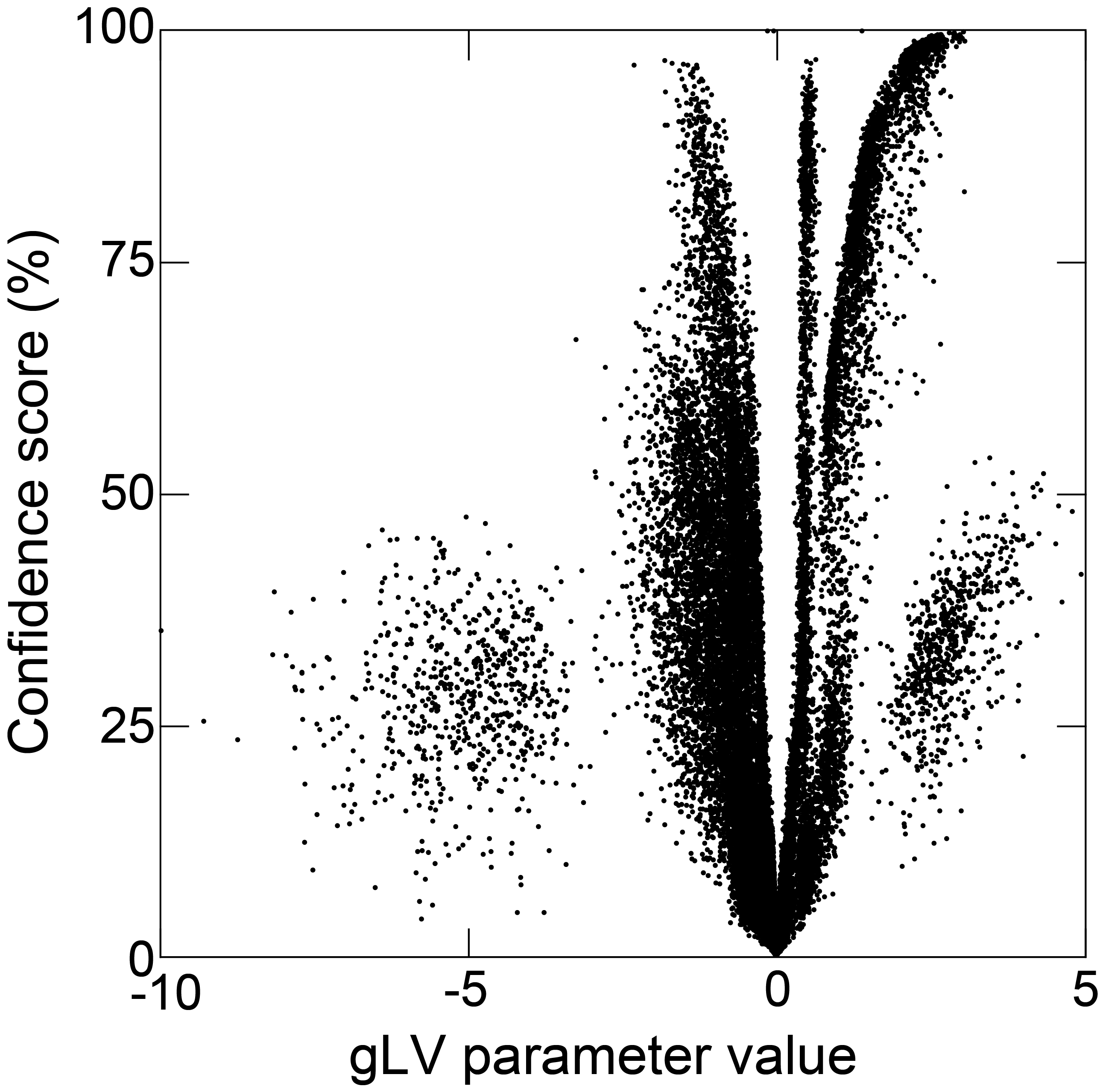

**Fig. S6** General proportionality between the absolute values of the generalized Lotka-Volterra (gLV) model parameters and their corresponding confidence scores. Each dot represents one parameter from one gLV model in the ensemble.

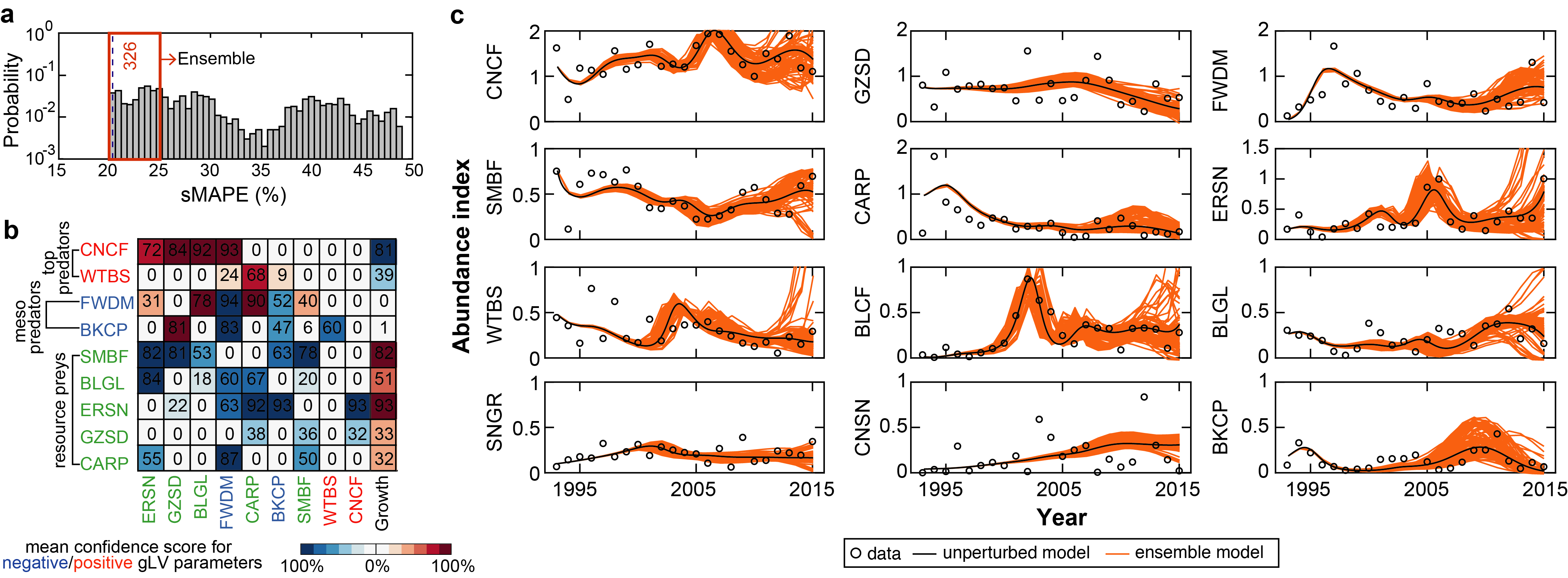

**Fig. S7** Modelling the Pool 26 fish community. **a** Probability distribution of fitting errors (sMAPE: symmetric mean absolute percentage error) among 1000 generalized Lotka-Volterra (gLV) models generated by perturbing the model given by our latent gradient regression (LGR) algorithm. The cutoff criteria (i.e., sMAPE ≤ 0.25) is the same as used for the La Grange pool and we obtained 326 models in the ensemble. **b** Ensemble-mean confidence scores for gLV parameters. The numbers in the square matrix made of the 9 rows and the first 9 columns are the mean confidence scores of pairwise interaction coefficients and indicate the likelihood that fish species on the column impacts fish species on the row. The numbers in the last column are the mean confidence scores of intrinsic growth rates and indicate the likelihood that population of each fish species grows (preys) or declines (predators) in the absence of the others. **c** Simulated trajectories of the 326 models in the ensemble. Unperturbed model is the best-fit model given by LGR.
